# Studies on CRMP2 SUMOylation-deficient transgenic mice reveal novel insights into the trafficking of NaV1.7 channels

**DOI:** 10.1101/2020.09.29.318774

**Authors:** Kimberly Gomez, Dongzhi Ran, Cynthia L. Madura, Aubin Moutal, Rajesh Khanna

## Abstract

Voltage-gated sodium channels are key players in neuronal excitability and pain signaling. Functional expression of the voltage-gated sodium channel NaV1.7 is under the control of SUMOylated collapsin response mediator protein 2 (CRMP2). If not SUMOylated, CRMP2 forms a complex with the endocytic proteins Numb, the epidermal growth factor receptor pathway substrate 15 (Eps15), and the E3 ubiquitin ligase Nedd4-2 to promote clathrin-mediated endocytosis of NaV1.7. We recently reported that CRMP2 SUMO-null knock-in (CRMP2^K374A/K374A^) female mice have reduced NaV1.7 membrane localization and currents in their sensory neurons. Preventing CRMP2 SUMOylation was sufficient to reverse mechanical allodynia in CRMP2^K374A/K374A^ female mice with neuropathic pain. Here we report that inhibiting clathrin assembly in nerve-injured male and female CRMP2^K374A/K374A^ mice, increased pain sensitivity in allodynia-resistant animals. Furthermore, Numb, Nedd4-2 and Eps15 expression was not modified in basal conditions in the dorsal root ganglia (DRG) of male and female CRMP2^K374A/K374A^ mice. Finally, silencing these proteins in DRG neurons from female CRMP2^K374A/K374A^ mice, restored the loss of sodium currents. Our study shows that the endocytic complex composed of Numb, Nedd4-2 and Eps15, is necessary for non SUMOylated CRMP2-mediated internalization of sodium channels *in vivo.*

## Introduction

NaV1.7 is a voltage-gated sodium channel highly expressed in nociceptive neurons and in the dorsal horn of the spinal cord (1). Biophysically, NaV1.7 manifests as a fast-activating and inactivating channel with a slow repriming (recovery from inactivation) kinetics, with sensitivity to tetrodotoxin (TTX-S) (2). NaV1.7 acts as a threshold channel to propagate action potentials in response to depolarizations of sensory neurons by noxious stimuli (3). Past research has established NaV1.7 as both necessary and sufficient for pain sensitivity (4, 5). Also well documented is the role of NaV1.7 during neuropathic and inflammatory pain, wherein NaV1.7 channel function and membrane expression are increased, resulting in hyperexcitability of dorsal root ganglia (DRG) neurons, likely via an amplification of subthreshold depolarizations (6).

Studies over the last decade in our laboratory have identified that collapsin response mediator protein 2 (CRMP2) as a regulator of diverse ion channels (7–14), including NaV1.7 (15–17). Regulation of NaV1.7 function follows CRMP2’s post-translational modification states, primarily phosphorylation and SUMOylation (i.e. addition of a small ubiquitin like modifier (SUMO)). In rodent models mimicking chronic neuropathic pain states, CRMP2 phosphorylation by cyclin-dependent kinase 5 (Cdk5) at Ser522 (S522) is increased (18–21); this is accompanied by enhanced SUMOylation by the E2-conjugating enzyme (Ubc9) at Lys374 (K374) (16, 22). Increased CRMP2 SUMOylation facilitates NaV1.7 membrane expression and increases the excitability of DRG neurons, and together these events may contribute to the expression of neuropathic pain (16, 17). Loss of CRMP2 SUMOylation results in: (i) reduced NaV1.7-CRMP2 binding; (ii) increased NaV1.7 internalization via association and recruitment of a tripartite complex of proteins containing the endocytic protein Numb, the E3 ubiquitin ligase neuronal precursor cell expressed developmentally downregulated-4 type 2 (Nedd4-2), and epidermal growth factor receptor pathway substrate 15 (Eps15); and (iii) decreased NaV1.7 surface expression and currents (16). This reduction can be rescued by blocking clathrin mediated endocytosis or by deleting the endocytic proteins Numb, Nedd4-2 or Eps15 (16). Numb is an endocytic adaptor protein that has been reported to bind to components of clathrin mediated endocytosis machinery (23). Numb can recruit Nedd4-2 to mark NaV1.7 for endocytosis by monoubiquitination (16, 24). Eps15 then binds to the monoubiquitinated channel to induce the initial curvature of the membrane and allow for the clathrin coated pit to form (25, 26).

To determine the role of CRMP2 SUMOylation *in vivo*, we generated CRMP2 K374A knock-in (CRMP2^K374A/K374A^) transgenic mice, in which the sole SUMOylation site on CRMP2 at Lysine 374 was replaced by an alanine (27). Although CRMP2 SUMOylation was abolished in both male and female homozygous mice, we unexpectedly discovered that CRMP2 dependent NaV1.7 trafficking was sexually dimorphic (28). In female mice only, germline loss of CRMP2 SUMOylation reduced NaV1.7–CRMP2 binding, NaV1.7 membrane localization and NaV1.7 currents in DRG sensory neurons, compared to their wildtype (WT) littermates (28). Inhibiting clathrin assembly with Pitstop2 (29), rescued the decreased sodium currents back to the levels observed in DRG from WT female mice. In contrast, none of these effects were observed in male mice, suggesting that the CRMP2^K374A/K374A^ mutation imposed a sex-specific regulation on NaV1.7. Of relevance to the role of NaV1.7 in pain, we found that CRMP2^K374A/K374A^ mice of both sexes failed to develop mechanical allodynia after a spinal nerve injury (SNI) (28), thus supporting the conclusion that CRMP2 SUMOylation-dependent regulation of NaV1.7 still holds in chronic neuropathic pain in male mice. Together, these studies underscore the critical role of CRMP2 SUMOylation as a regulatory mechanism underlying NaV1.7 functional expression.

In this study, we asked two questions. First, is the lack of mechanical allodynia in male CRMP2^K374A/K374A^ mice due to increased internalization of NaV1.7? And second, is the decrease in NaV1.7 currents in female CRMP2^K374A/K374A^ mice rely on the endocytic proteins Numb, Nedd4-2 and Eps15? Here, using CRMP2^K374A/K374A^ mice, we show that: (i) inhibition of clathrin-mediated endocytosis restores nociception following SNI in male mice; (ii) Numb, Nedd4-2 and Eps15 expression is no different between DRGs from males and female mice; and (iii) knocking down these endocytic proteins in DRG neurons restores the loss of sodium currents observed in female CRMP2^K374A/K374A^ mice. Together, our data resolve the mechanism of decreased NaV1.7 membrane localization, currents and lack of allodynia in male and female CRMP2^K374A/K374A^ mice.

## Results

### In vivo inhibition of clathrin assembly restores mechanical allodynia in male CRMP2^K374A/K374A^ mice with spinal nerve injury

Our previous in vitro studies found that the CRMP2^K374A/K374A^ genotype leads to a reduction of sodium current in female, but not male, mice (28). Inhibiting clathrin assembly with the small molecule Pitstop2 (16), rescued the decrease in sodium currents observed in DRG neurons from female CRMP2^K374A/K374A^ mice (28). Subsequent *In vivo* studies revealed that, following SNI, both CRMP2^K374A/K374A^ male and female mice become resistant to the development of mechanical allodynia compared to their WT littermates (28). From these data, one can surmise that although there is no impact of loss of CRMP2 SUMOylation on sodium currents in male mice, inflicting a neuropathic pain injury uncovers an effect in both genders (28). To resolve this discrepancy, we hypothesized that increased internalization of NaV1.7 in male CRMP2^K374A/K374A^ mice compromises its role in neuropathic pain. As inhibiting clathrin assembly normalized NaV1.7 current loss in female CRMP2^K374A/K374A^ mice, we asked if Pitstop2 could precipitate mechanical allodynia in male CRMP2^K374A/K374A^ mice with neuropathic pain. In male CRMP2^K374A/K374A^ mice, we found higher paw withdrawal thresholds at 42 days after in mice subjected to SNI compared to their WT littermates (Fig. 1). After intrathecal administration of 0.1% DMSO (vehicle), WT mice maintained a low paw withdrawal threshold, whilst CRMP2^K374A/K374A^ mice showed a persistent resistance to mechanical allodynia (Fig. 1). In contrast, administration of 2 μg/5 μl of Pitstop2 significantly decreased the paw withdrawal thresholds in male CRMP2^K374A/K374A^ mice 1 hour following the injection and then gradually recovered. No significant changes were observed in WT male mice (Fig. 1). Taken together, this data suggests that, in male CRMP2^K374A/K374A^ mice, enhanced clathrin mediated endocytosis contributes to the resistance to mechanical allodynia in CRMP2^K374A/K374A^ mice with SNI. This is consistent with our previous observations of increased CRMP2 SUMOylation in male rats with SNI (17).

**Figure 1.**
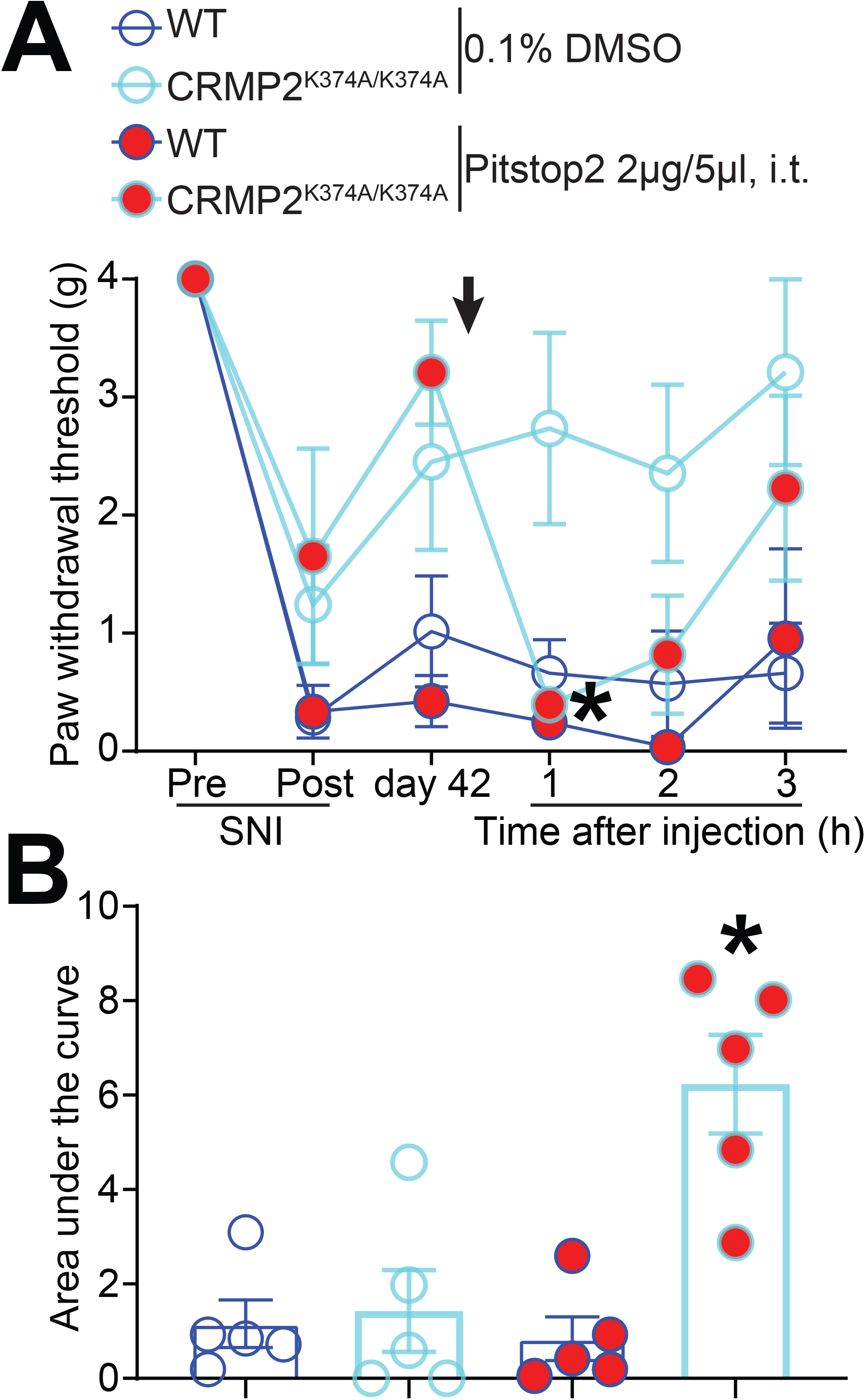
Pitstop2 rescues mechanical allodynia in male CRMP2^K374A/K374A^ knock-in mice with a spared nerve injury (SNI). Paw withdrawal thresholds of age-matched and genotyped WT and CRMP2^K374A/K374A^ mice were measured at baseline and for six weeks following SNI. Post SNI, von Frey testing was confined to the sural nerve innervated region of the paw. (**A**) Time course showing that, at 42 days following SNI, male CRMP2^K374A/K374A^ mice do not develop mechanical allodynia, which can however be precipitated by Pitstop2 administered by a lumbar puncture (2 μg in 5 μl) and followed over 3 hours. *p<0.05 two-way ANOVA with Sidak’s post hoc test. (**B**) Area under the curve for paw withdrawal thresholds was derived using the trapezoid method. Error bars indicate mean ± SEM. *p<0.05 Kruskal-Wallis test. The experiments were conducted by an investigator blinded to the genotype and treatment (n=5 each).

### Silencing Numb, Nedd4-2 and Eps15 rescues the decrement of sodium currents in female CRMP2^K374A/K374A^ mice

Our previous study established that non-SUMOylated CRMP2 recruits Numb, Nedd4-2 and Eps15 to regulate internalization of NaV1.7 channels (16). In basal conditions, in male and female CRMP2^K374A/K374A^ mice DRG, NaV1.7 currents contribute to ~58% and ~76% of the total sodium currents, respectively (28). However, Na^+^ currents were decreased only in female CRMP2^K374A/K374A^ mice while we could not measure any difference in their male littermates. We ruled out that the failure to observe this loss of NaV1.7 currents in male CRMP2^K374A/K374A^ DRGs was not due to lack of the neurotrophic factors nerve growth factor (NGF) or brain derived neurotrophic factor (BDNF) nor due to increased basal endocytic activity (28). However, our previous work did not determine the level of expression of the endocytic proteins Numb, Nedd4-2 and Eps15. Consequently, here we asked if the levels of these proteins could explain the sex specific outcomes of CRMP2^K374A/K374A^ mice. We dissected lumbar DRGs from male and female WT mice and detected the level of expression of Numb, Nedd4-2 and Eps15 by western blot. We did not find any significant difference of expression of these endocytic proteins between male and female animals (Fig. 2). Hence, the sex difference in NaV1.7 currents cannot not be attributed to changes in the expression of endocytic proteins that regulate its internalization.

**Figure 2.**
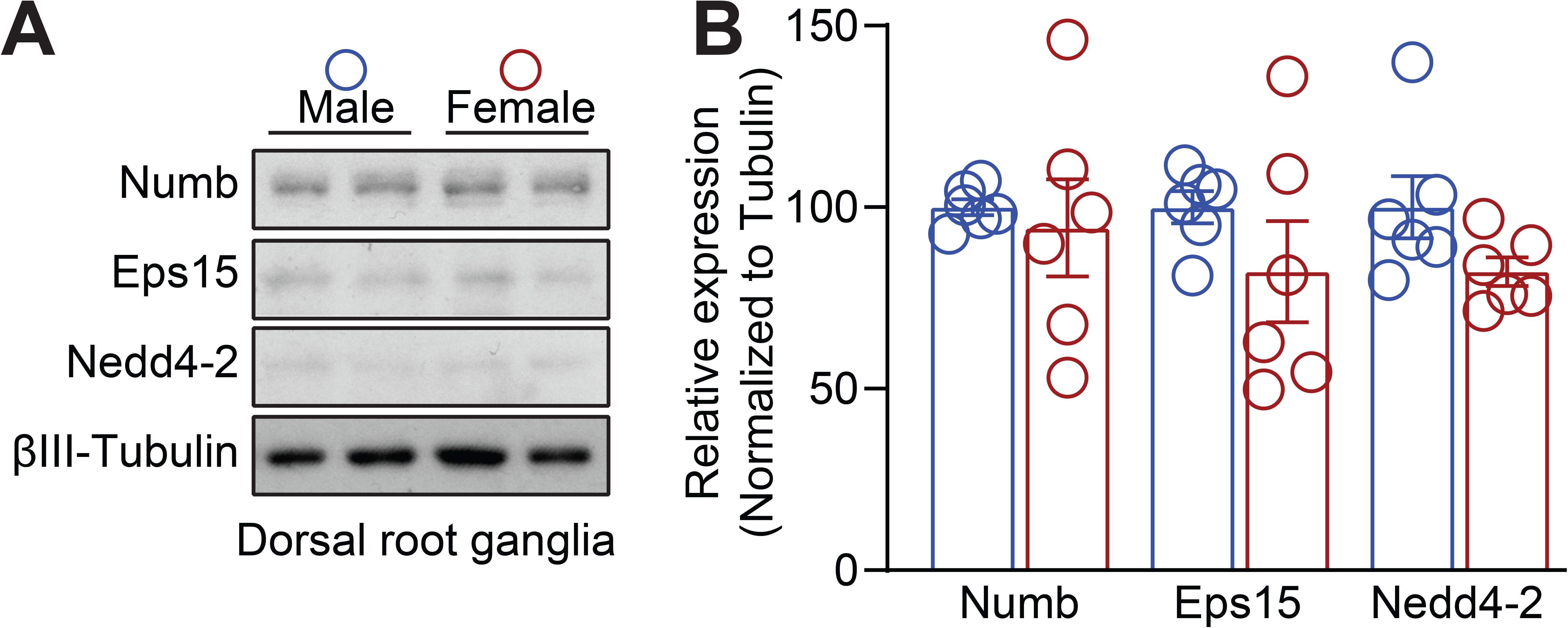
Expression of Numb, Nedd4-2 and Eps15 in male and female DRGs. **A**) Representative Western blots showing the expression of the endocytic proteins Numb, Nedd4-2 and Eps15 in lumbar DRG from male and female wildtype mice. βIII-tubulin served as a loading control. (**B**) Bar graph with scatter plot showing no difference of expression of the proteisn between male and female DRG (n = 6).

Having demonstrated that inhibiting clathrin assembly in CRMP2^K374A/K374A^ mice with neuropathic pain restores mechanical sensitivity, we next investigated the functional consequences of knocking-down each of the proteins that constitute the endocytic machinery. We used specific siRNAs (validated in (16)) to knockdown expression of Numb (Fig. 3), Nedd4-2 (Fig. 4), or Eps15 (Fig. 5) DRG neurons. Figures 3A, 4A and 5A display representative sodium currents recorded from female WT and CRMP2^K374A/K374A^ DRG neurons transfected with siRNA control and siRNA against Numb (Fig. 3A), Nedd4-2 (Fig. 4A), or Eps15 (Fig. 5A). Silencing Numb, Nedd4-2 and Eps15 in CRMP2^K374A/K374A^, restored the decreased sodium currents density (106.5%, 238.9% and 279.6%, respectively), back to the levels observed in DRG from CRMP2^K374A/K374A^ transfected with a control siRNA (Fig. 3B, 4B and 5B). In contrast, silencing these proteins in DRG from WT mice had no effect (Fig. 3B, 4B and 5B). To account for the heterogeneity of the DRG population in terms of neuronal size, peaks were normalized by cell capacitance and subsequently displayed as peak current density (pA/pF). Again, there was restoration of peak currents in CRMP2^K374A/K374A^ mice after knocking-down the endocytic proteins, while in WT mice there were no differences noted (Figs. 3C, 4C and 5C). To test if silencing Numb, Nedd4-2 and Eps15 could cause changes in channel gating, we next calculated the voltage-dependent activation and inactivation properties of sodium currents in DRG neurons. Comparing the midpoint potentials (V_1/2_) and slope factors (*k*) in response to changes in command voltages (Table 1) of whole-cell ionic conductance, allowed us to measure changes in activation and inactivation of sodium currents for the DRG neurons transfected with siRNAs. Representative Boltzmann fits are shown in Figures 3D, 4D and 5D. There were no significant differences in the steady-state activation and inactivation properties of sodium currents, per analysis of *V_1/2_* and *k* values, between DRG neurons of any condition (Table 1). Altogether, our results show that the lack of any of these endocytic proteins is sufficient to disrupt the endocytic complex to affect a rescue of the decreased sodium currents. This is entirely consistent with past results from rat DRG neurons overexpression of a sumo null CRMP2 (16).

**Figure 3.**
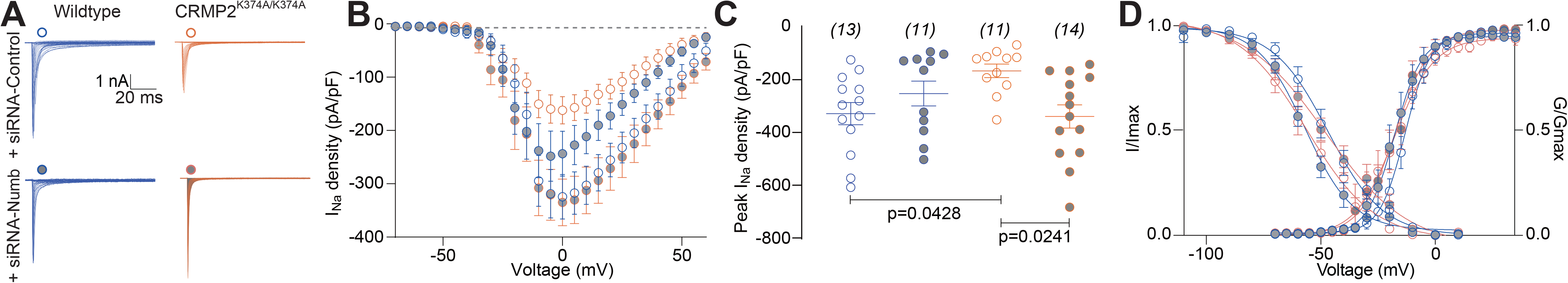
CRMP2-mediated decrease in sodium currents observed in DRGs from CRMP2^K374A/K374A^ mice is rescued in cells lacking Numb. (**A**) Representative sodium current traces recorded from small-sized DRG neurons of WT and CRMP2^K374A/K374A^ female mice, in response to depolarization steps from −70 to +60 mV from a holding potential of −60 mV. (**B**) Summary of current-voltage curves and (**C**) peak currents (pA/pF) from DRG neurons. Total sodium current density was significantly smaller in DRGs from homozygous mice vs. DRG neurons from WT+siRNA-Control (n=11-15 cells/condition, p = 0.0428, Kruskal-Wallis test with Dunn’s post hoc), and vs. DRG neurons from CRMP2^K374A/K374A^+siRNA-Numb (n=11-15, p = 0.0241, Kruskal-Wallis test with Dunn’s post hoc). (**D**) Boltzmann fits for normalized conductance *G/Gmax* voltage relations for voltage dependent activation and inactivation of the sensory neurons. Half-maximal activation and inactivation (*V*_*1/2*_) and slope values (*k*) for activation and inactivation are presented in Table 1. There were no significant differences in *V*_*1/2*_ and *k* values of activation and inactivation between genotype and treatment (One-way ANOVA with Dunnett’s post hoc test). Error bars indicate mean ± SEM.

**Figure 4.**
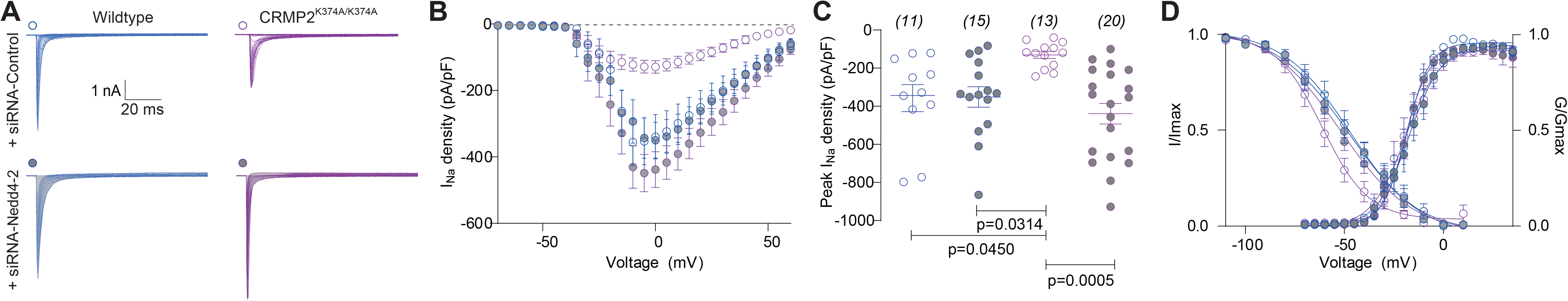
CRMP2-mediated decrease in sodium currents observed in DRGs from CRMP2^K374A/K374A^ mice is rescued in cells lacking Nedd4-2. (**A**) Representative sodium current traces recorded from small-sized DRG neurons of WT and CRMP2^K374A/K374A^ female mice, in response to depolarization steps from −70 to +60 mV from a holding potential of −60 mV. (**B**) Summary of current-voltage curves and (**C**) peak currents (pA/pF) from DRG neurons. Total sodium current density was significantly smaller in DRGs from homozygous female mice vs. DRG neurons from WT+siRNA-Control (n=12-12 cells/condition, p = 0.0450, Kruskal-Wallis test with Dunn’s post hoc), vs. DRG neurons from WT+siRNA-Nedd4-2 (n=12-15, p = 0.0314, Kruskal-Wallis test with Dunn’s post hoc), and vs. DRG neurons from CRMP2^K374A/K374A^+siRNA-Nedd4-2 (n=12-19, p = 0.0005, Kruskal-Wallis test with Dunn’s post hoc). (**D**) Boltzmann fits for normalized conductance *G/Gmax* voltage relations for voltage dependent activation and inactivation of the sensory neurons. Half-maximal activation and inactivation (*V_1/2_*) and slope values (*k*) for activation and inactivation are presented in Table 1. There were no significant differences in *V_1/2_* and *k* values of activation and inactivation between genotype and treatment (One-way ANOVA with Dunnett’s post hoc test). Error bars indicate mean ± SEM.

**Figure 5.**
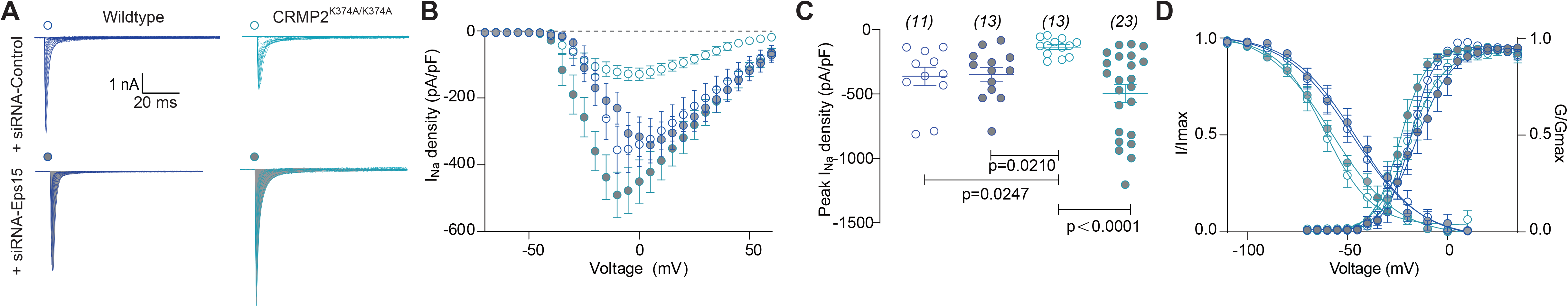
CRMP2-mediated decrease in sodium currents observed in DRGs from CRMP2^K374A/K374A^ mice is rescued in cells lacking Eps15. (**A**) Representative sodium current traces recorded from small-sized DRG neurons of WT and CRMP2^K374A/K374A^ female mice, in response to depolarization steps from −70 to +60 mV from a holding potential of −60 mV. (B) Summary of current-voltage curves and (C) peak currents (pA/pF) from DRG neurons. Total sodium current density was significantly smaller in DRGs from homozygous female mice vs. DRG neurons from WT+siRNA-Control (n=12-12 cells/condition, p = 0.0247, Kruskal-Wallis test with Dunn’s post hoc), vs. DRG neurons from WT+siRNA-Eps15 (n=12-13, p = 0.0210, Kruskal-Wallis test with Dunn’s post hoc), and vs. DRG neurons from CRMP2^K374A/K374A^+siRNA-Eps15 (n=12-17, p < 0.0001, Kruskal-Wallis test with Dunn’s post hoc). (**D**) Boltzmann fits for normalized conductance *G/Gmax* voltage relations for voltage dependent activation and inactivation of the sensory neurons. Half-maximal activation and inactivation (*V_1/2_*) and slope values (*k*) for activation and inactivation are presented in Table 1. There were no significant differences in *V_1/2_* and *k* values of activation and inactivation between genotype and treatment (One-way ANOVA with Dunnett’s post hoc test). Error bars indicate mean ± SEM.

**Table 1.**
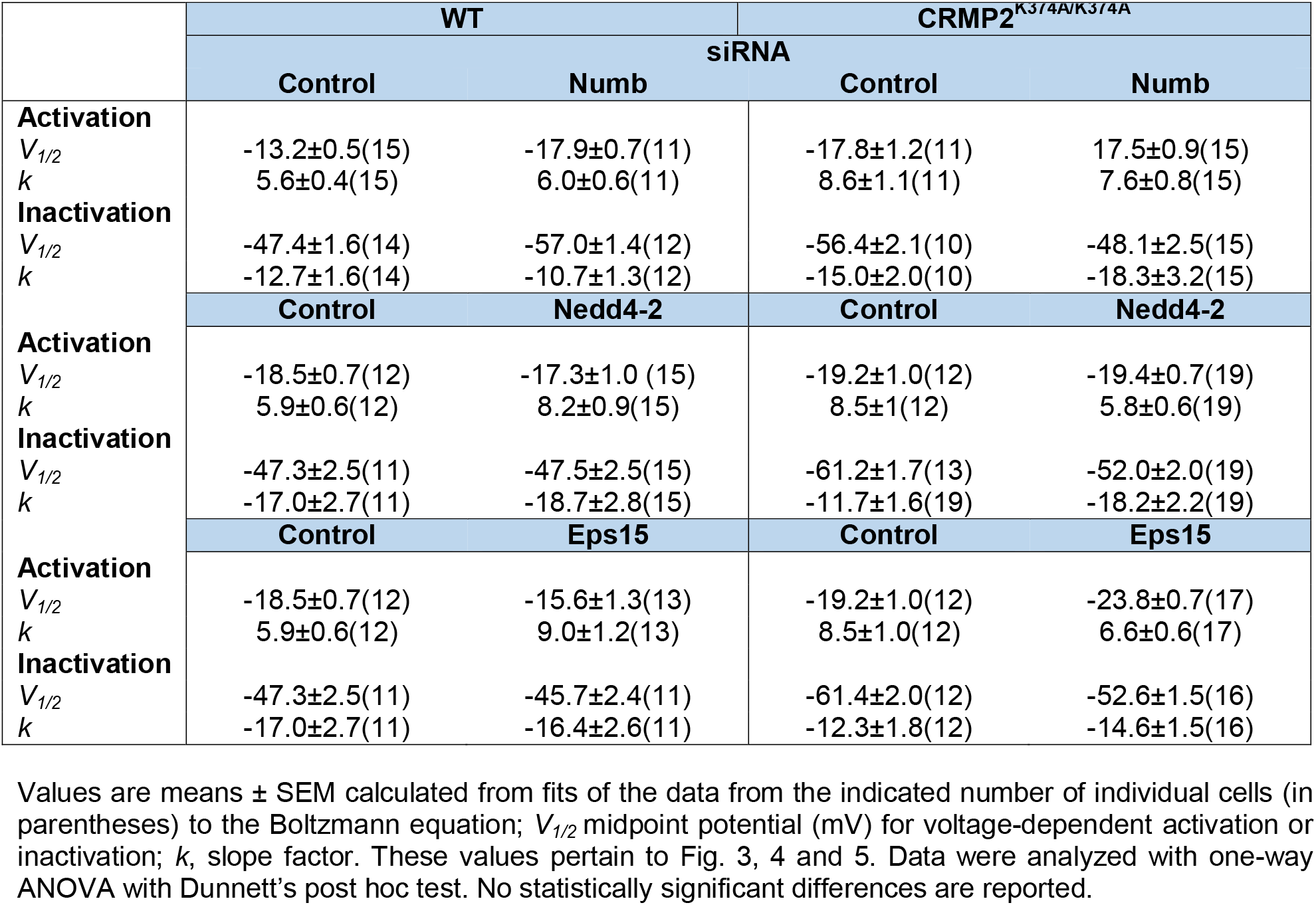
Effects of silencing Numb, Nedd4-2 and Eps15 on gating properties of sodium channels in DRG neurons from WT and CRMP2K374A/K374A female mice.

|  | WT |  | CRMP2<sup>K374A/K374A</sup> |  |
| --- | --- | --- | --- | --- |
|  | siRNA |  |  |  |
|  | Control | Numb | Control | Numb |
| **Activation** |  |  |  |  |
| $V_{1/2}$ | -13.2±0.5(15) | -17.9±0.7(11) | -17.8±1.2(11) | 17.5±0.9(15) |
| $k$ | 5.6±0.4(15) | 6.0±0.6(11) | 8.6±1.1(11) | 7.6±0.8(15) |
| **Inactivation** |  |  |  |  |
| $V_{1/2}$ | -47.4±1.6(14) | -57.0±1.4(12) | -56.4±2.1(10) | -48.1±2.5(15) |
| $k$ | -12.7±1.6(14) | -10.7±1.3(12) | -15.0±2.0(10) | -18.3±3.2(15) |
|  | Control | Nedd4-2 | Control | Nedd4-2 |
| **Activation** |  |  |  |  |
| $V_{1/2}$ | -18.5±0.7(12) | -17.3±1.0 (15) | -19.2±1.0(12) | -19.4±0.7(19) |
| $k$ | 5.9±0.6(12) | 8.2±0.9(15) | 8.5±1(12) | 5.8±0.6(19) |
| **Inactivation** |  |  |  |  |
| $V_{1/2}$ | -47.3±2.5(11) | -47.5±2.5(15) | -61.2±1.7(13) | -52.0±2.0(19) |
| $k$ | -17.0±2.7(11) | -18.7±2.8(15) | -11.7±1.6(19) | -18.2±2.2(19) |
|  | Control | Eps15 | Control | Eps15 |
| **Activation** |  |  |  |  |
| $V_{1/2}$ | -18.5±0.7(12) | -15.6±1.3(13) | -19.2±1.0(12) | -23.8±0.7(17) |
| $k$ | 5.9±0.6(12) | 9.0±1.2(13) | 8.5±1.0(12) | 6.6±0.6(17) |
| **Inactivation** |  |  |  |  |
| $V_{1/2}$ | -47.3±2.5(11) | -45.7±2.4(11) | -61.4±2.0(12) | -52.6±1.5(16) |
| $k$ | -17.0±2.7(11) | -16.4±2.6(11) | -12.3±1.8(12) | -14.6±1.5(16) |

Values are means ± SEM calculated from fits of the data from the indicated number of individual cells (in parentheses) to the Boltzmann equation; $V_{1/2}$ midpoint potential (mV) for voltage-dependent activation or inactivation; $k$ , slope factor. These values pertain to Fig. 3, 4 and 5. Data were analyzed with one-way ANOVA with Dunnett's post hoc test. No statistically significant differences are reported.

## Discussion

Our results provide further mechanistic insights into how loss of CRMP2 SUMOylation decreases Nav1.7 currents in female mice and mechanical allodynia in male mice. We demonstrate that inhibiting clathrin assembly in male CRMP2^K374A/K374A^ mice, following an injury that results in neuropathic pain, restores mechanical sensitivity. When determining the expression levels of the CRMP2 interacting endocytic proteins in the DRG, we found no differences between sexes under basal conditions. Furthermore, the reduction of Na^+^ currents in DRG neurons from female CRMP2^K374A/K374A^ mice were normalized by silencing Numb, Nedd4-2 and Eps15.

The dynamic process of SUMOylation/deSUMOylation of CRMP2 controls the membrane surface expression and current density of NaV1.7 channels in rodent and human DRGs (15, 16). We have demonstrated that this CRMP2-regulation is selective for NaV1.7, and therefore spares the other voltage-gated Na^+^ channels in DRG neurons (16). In chronic neuropathic pain, we found that enhanced NaV1.7 functional expression correlates with an increase of CRMP2 SUMOylation (16, 30). Interestingly, expressing a SUMO-null CRMP2 mutant (K374A) in rats with neuropathic pain was sufficient to reverse mechanical allodynia [9]. Also, CRMP2^K374A/K374A^ male and female mice are resistant to the development of mechanical allodynia after a spared nerve injury (SNI) [19] (Figs. 1 and 6). Because NaV1.7 endocytosis is clathrin-dependent (16), we reasoned that inhibiting clathrin assembly with Pitstop 2 in vivo may restore mechanical allodynia in these animals. We show that this resistance of CRMP2^K374A/K374A^ mice is dependent on clathrin-mediated endocytosis (Figs. 1 and 6). In other words, Pitstop 2 contributes to the re-establishment of mechanical allodynia in these mice probably by maintaining NaV1.7 membrane expression where it is available for voltage-dependent activation. This finding correlates with our previous reports that Pitstop 2 prevents Na^+^ current density reductions imposed by loss of CRMP2 SUMOylation (16, 28) (Fig. 6B). Therefore, our data suggests that the association of non-SUMOylated CRMP2 with the clathrin machinery contributes to mitigate pain. This in accordance with our observations that interfering with SUMOylation of CRMP2, promotes clathrin-mediated endocytosis of NaV1.7 (16) and reverses mechanical allodynia (30).

**Figure 6.**
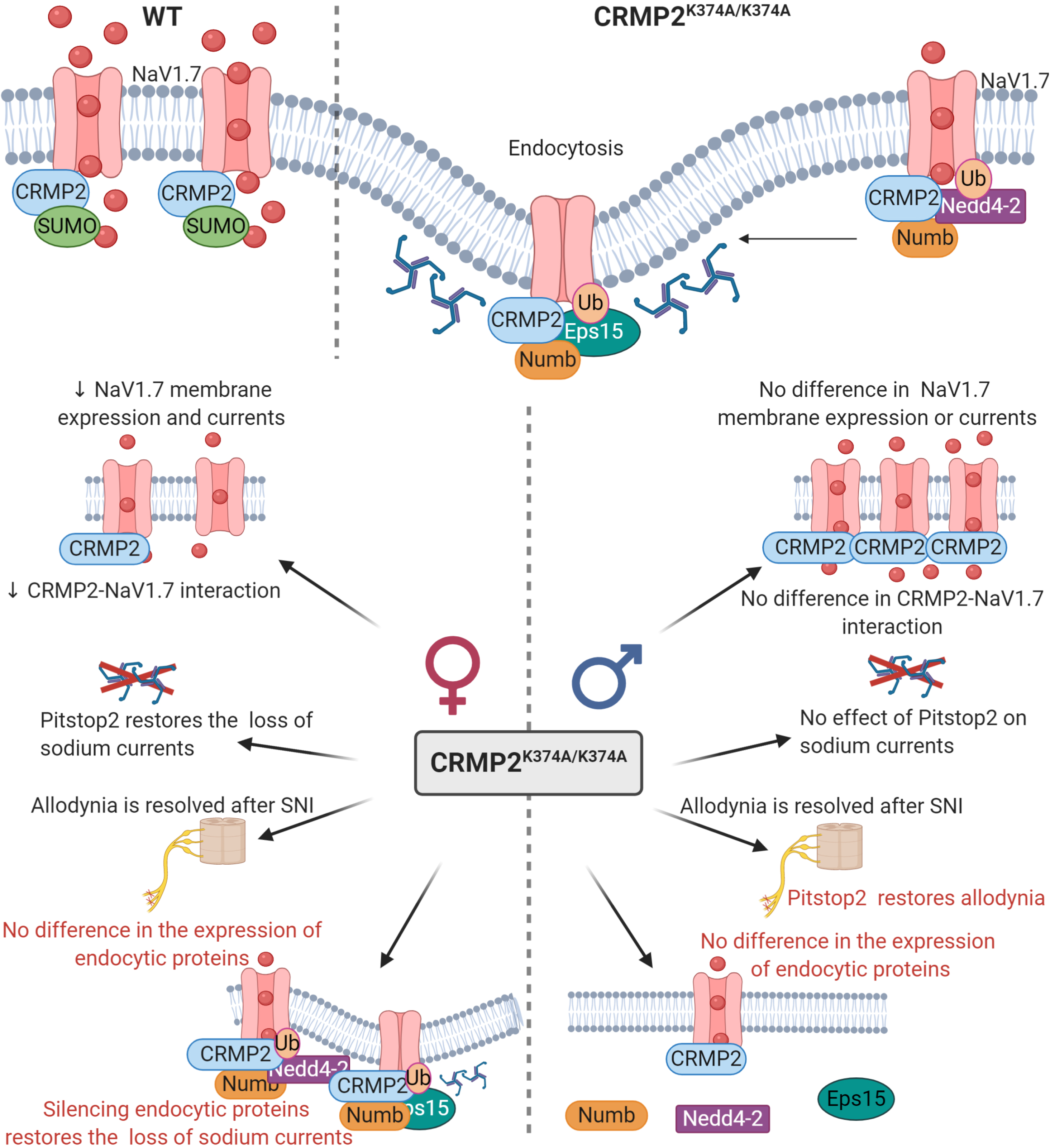
Schematic representation of CRMP2 regulation of NaV1.7 channels. *Top:* In wildtype (WT) mice (*left*), when CRMP2 is SUMOylated it interacts with NaV1.7 channels to enhance their trafficking to the plasma membrane. In CRMP2^K374A/K374A^ mice (*right*), nonSUMOylated CRMP2 recruits Numb, Nedd4-2 and Eps15 to trigger clathrin-mediated endocytosis of NaV1.7 channels. *Bottom:* Sex specific differences of female (*left*) and male (*right*) CRMP2^K374A/K374A^ mice compared to their WT littermates. In female mice: (i) NaV1.7 membrane expression, currents, and CRMP2-NaV1.7 interaction are reduced (28), (ii) Pitstop2 restores the loss of sodium currents and allodynia (28), (iii) in basal conditions no difference in the expression of endocytic proteins is seen and, (iv) silencing Numb, Nedd4-2 and Eps15 restores the loss of sodium currents. In male mice: (i) NaV1.7 membrane expression, currents, and CRMP2-NaV1.7 interaction are not modified (28), (ii) Pitstop2 has no effect on sodium currents (28) but restores mechanical allodynia, and (iii) in basal conditions no difference in the expression of Numb, Nedd4-2 and Eps15 is observed. Red text represents the data obtained in this study.

Loss of CRMP2 SUMOylation increases its association with Numb, Nedd4-2 and Eps15, and accumulates NaV1.7 channels into rab5 positive early recycling endosomes (16). Numb is involved in clathrin-dependent endocytosis at the plasma membrane; it associates with the appendage domain of α adaptin, a subunit of the clathrin adaptor complex AP2, a major component of clathrin-coated pits (23). Numb can interact *in vivo* and *in vitro* with Eps15, a component of the endocytic machinery (31), that also interacts with AP2 (32). The E3 ubiquitin ligase Nedd4-2 is a potent post translational regulator of NaV1.7 and NaV1.8 voltage-gated sodium channels (24). Its downregulation leads to hyperexcitability of DRG neurons and contributes to the onset of chronic neuropathic pain (24). We recently reported that ~76% of the sodium current is contributed by NaV1.7 sodium in DRGs from WT and CRMP2^K374A/K374A^ female mice (28). This fraction was determined by applying PF05089771, a NaV1.7-specific blocker (33). In their male counterparts, NaV1.7 sodium currents represented 53% and 58% of total sodium currents from WT and CRMP2^K374A/K374A^ DRGs, respectively [19]. To assess if this difference between males and females could be due to differences in the expression of the proteins that regulate NaV1.7 endocytosis, we performed western blot analyses. In naïve wildtype mice, the protein levels of Numb, Nedd4-2 and Eps15 in the lumbar DRG are similar between males and females (Fig. 2 and 6B). These data leave open the possibility (not tested here) that in females there is a lesser degree of interaction between CRMP2 and these endocytic proteins in the DRG, compared to males. In support of this possibility, we found that CRMP2 interaction with NaV1.7 was marginal in wildtype males compared to females (28).

CRMP2^K374A/K374A^ mutation decreases sodium currents in a sex-specific manner (28). Total sodium currents are decreased (~40%) in female CRMP2^K374A/K374A^ DRG but not in males [19] (Fig. 6B). For this reason, we decided to determine the functional relevance of these endocytic proteins in our female knock-in mice. Recordings of total sodium currents revealed that the decreased current density, ranging from 50 to ~63.7%, was prevented when Numb, Nedd4-2 and Eps15 were individually silenced in acutely dissociated DRG neurons (Figs. 3, 4 and 5). By knocking down each of these proteins individually in sensory neurons, we showed that they coordinate NaV1.7 internalization when CRMP2 SUMOylation is lost. These results correlate with our finding that following specific reduction of these CRMP2-interacting proteins, the inhibition of NaV1.7 currents by loss of CRMP2 SUMOylation is rescued (16).

In summary, in WT mice, SUMOylation of CRMP2 allows for CRMP2-NaV1.7 interaction and NaV1.7 channel peregrination to the plasma membrane. In CRMP2^K374A/K374A^ mice, non-SUMOylated CRMP2 triggers NaV1.7 endocytosis by recruiting Numb, Nedd4-2 which ubiquitinates Nav1.7 (24) and Eps15 which induces membrane curvature (26) (Fig. 6). However, while the above mechanism is true in female, but not male, DRG neurons from naïve mice, we found that in neuropathic pain conditions, pain assessment in male and female animals is indistinguishable (28) (Fig. 1). Comparing female CRMP2^K374A/K374A^ with WT mice, we note: (i) NaV1.7 membrane expression, currents, and CRMP2-NaV1.7 interaction are decreased, (ii) Pitstop2 restores the loss of sodium currents and mechanical allodynia, (iii) no difference in the expression of endocytic proteins is observed in basal conditions, and (iv) silencing these endocytic proteins restores the loss of sodium currents. This granularity in regulation of NaV1.7 membrane expression, a ‘coding’ of the function of the channel, may be utilized to design new therapeutics for chronic pain. In support of this, we used a CRMP2 SUMOylation blocking peptide strategy as a proof of concept to demonstrate that preventing this modification can reverse established chronic allodynia in neuropathic pain (34). This unique approach will allow for the inhibition of enhanced NaV1.7 function reported in neuropathic pain while leaving intact the pool of the channels participating in physiological pain sensation.

## Materials and methods

### Animals

All animal use was conducted in accordance with the National Institutes of Health guidelines, and the study was carried out in strict accordance with recommendations in the Guide for the Care and Use of Laboratory Animals of the University of Arizona (Protocol #: 16-141). Wildtype and CRMP2^K374A/K374A^ mice were housed and bred in the University of Arizona Laboratory Animal Research Center. Mice were housed in groups of 4-5 in a dedicated housing facility with ad libitum access to food and water on a 12-hour light/dark cycle. All animal procedures were performed in accordance with the policies and recommendations of the National Institutes of Health, and with approval from the animal care and use committee of the University of Arizona for the handling and use of laboratory animals.

### Antibodies and siRNAs

Biochemical analysis of protein content by Western blot used the following antibodies: Numb (cat. no. ab4147; Abcam), Eps15 (cat. no. ab174291; Abcam), Nedd4-2 (cat. no. ab131167 RRID: AB_11157800; Abcam), and βIII-Tubulin (cat. no. G712A; Promega). For RNA interference, siRNA Numb (5’-TAACTGGGAAGCTACACTTTCCAGT-3’), siRNA Nedd4-2 (5’-CATACTATGTCAATCATAATT-3’), siRNA Eps15 (5’-CCCAGGCAATGATAGTCCCAAAGAA-3’), and siRNA control (cat. no. 12935300) were obtained from Thermo Fisher Scientific.

### Western blot preparation and analysis

Indicated samples were loaded on 4–20% Novex gels (cat. no. EC60285BOX; Thermo Fisher Scientific). Proteins were transferred for 1 h at 120 V using TGS [25 mM Tris, pH 8.5,192 mM glycine, 0.1% (mass/vol) SDS], 20% (vol/vol) methanol as transfer buffer to PVDF membranes (0.45 μm; cat. no. IPVH00010; Millipore), preactivated in pure methanol. After transfer, the membranes were blocked at room temperature for 1 h with TBST (50 mM Tris·HCl, pH 7.4, 150 mM NaCl, 0.1% Tween 20) with 5% (mass/vol) nonfat dry milk, and then incubated separately in indicated primary antibodies in TBST, 5% (mass/vol) BSA, overnight at 4 °C. Following incubation in HRP-conjugated secondary antibodies from Jackson Immuno Research, blots were revealed by enhanced luminescence (WBKLS0500; Millipore) before exposure to photographic film. Films were scanned, digitized, and quantified by using Un-Scan-It gel scanning software (version 7.1; Silk Scientific).

### Preparation of acutely dissociated dorsal root ganglia neurons from wildtype (WT) and CRMP2^K374A/K374A^ mice

Wildtype and CRMP2^K374A/K374A^ mice were deeply anaesthetized with isoflurane overdose (5% in oxygen) and sacrificed by decapitation. Dorsal root ganglia (DRG) were quickly removed, trimmed at their roots, and enzymatically digested in 3 mL bicarbonate-free, serum-free, sterile DMEM (Cat# 11965, Thermo Fisher Scientific, Waltham, MA) solution containing neutral protease (1.87 mg/ml, Cat#LS02104; Worthington, Lakewood, NJ) and collagenase type I (3 mg/mL, Cat# LS004194, Worthington, Lakewood, NJ) and incubated for 50 minutes at 37°C under gentle agitation. Dissociated DRG neurons were then gently centrifuged to collect cells and washed with DRG media (DMEM containing 1% penicillin/streptomycin sulfate from 10,000 μg/mL stock, and 10% fetal bovine serum (Hyclone). Collected cells were resuspended in Nucleofector transfection reagent containing siRNA at a working concentration of 600 nM. Then, cells were subjected to electroporation protocol O-003 in an Amaxa Biosystem (Lonza) and plated onto 12-mm poly-D-lysine- and laminin-coated glass coverslips. Transfection efficiencies were routinely between 20% and 30%, with approximately ∼10% cell death. siRNA transfection was verified by GFP fluorescence.

### Whole-cell electrophysiological recordings of sodium currents in acutely dissociated DRG neurons from WT and CRMP2^K374A/K374A^ mice

All recordings were obtained from acutely dissociated DRG neurons from WT and CRMP2^K374A/K374A^ mice, using procedures adapted from those as described in our previously published study (28). Whole-cell voltage-clamp recordings were performed between 48 h and 72 h after transfection at room temperature, using an EPC 10 Amplifier-HEKA. The external solution consisted of (in mM): 140 NaCl, 30 tetraethylammonium chloride, 10 D-glucose, 3 KCl, 1 CaCl2, 0.5 CdCl2, 1 MgCl2, and 10 HEPES (pH 7.3, mOsm/L = 310-315) and internal solution contained the following (in mM): 140 CsF, 10 NaCl, 1.1Cs-EGTA, and 15 HEPES (pH 7.3, mOsm/L = 290-310). DRG neurons were subjected to current-voltage (I-V) and activation/inactivation voltage protocols as follows: (a) I-V protocol: from a −60 mV holding potential, cells were depolarized in 150-millisecond voltage steps from −70 to +60 mV (5-mV increments) which allowed acquisition of current density values such that we could analyze activation of sodium channels as a function of current vs voltage and infer peak current density (normalized to cell capacitance (in picofarads, pF)), which occurred between ~0 to 10 mV; (b) inactivation protocol: from a holding potential of −60 mV, cells were subjected to hyperpolarizing/repolarizing pulses for 1 second between −120 to 10 mV (+10 mV steps). Pipettes were pulled from standard wall borosilicate glass capillaries (Warner Instruments) with a horizontal puller (Model P-97, Sutter Instruments) and heat-polished to final resistances of 1–4 MΩ when filled with internal solutions. Whole-cell capacitance and series resistance of electrode were compensated. Linear leak currents were digitally subtracted through P/4 method for voltage clamp experiments. Signals were filtered at 10 kHz and digitized at 10–20 kHz. Cells in which series resistance was more than 15 MΩ over the course of an experiment were omitted from datasets. Analysis was performed by using Fitmaster software (HEKA) and Origin 9.0 software (OriginLab).

### Spared nerve injury (SNI) model of neuropathic pain

Age-matched and genotyped WT and CRMP2^K374A/K374A^ mice were habituated in plastic chambers on a mesh floor. Calibrated von Frey filaments with sequentially increasing spring coefficients were applied to the hind paw of each mouse, which allowed for the consistent application of constant force stimuli. The test was performed on the lateral part of the right plantar surface where the sural nerve innervates the hind paw. One filament was applied 5 times in a round of testing. The filament force evoking paw withdrawal more than 3 times in a round of testing was defined as the mechanical threshold. The cutoff threshold was 4 g. For the spared nerve injury (SNI) model, we transected the common peroneal and tibial branches of the right sciatic nerve with ~1 mm of nerve removed and left the sural nerve intact.

### Statistical analyses

All data was first tested for a Gaussian distribution using a D’Agostino-Pearson test (Prism 8 Software, Graphpad, San Diego, CA). The statistical significance of differences between means was determined by a parametric ANOVA followed by Dunnett’s post hoc or a non-parametric Kruskal Wallis test followed by Dunn’s post-hoc test depending on if datasets achieved normality. Behavioral data with a time course were analyzed by Two-way ANOVA with Sidak’s post hoc test. Differences were considered significant if p≤ 0.05. Error bars in the graphs represent mean ± SEM. All data were plotted in Prism 8.

## Abbreviations

Cdk5: Cyclin-dependent kinase 5
CRMP2: Collapsin response mediator protein 2
DRG: Dorsal root ganglia
Eps15: Epidermal growth factor receptor pathway substrate 15
k: Slope factor
NaV1.7: voltage-gated sodium channel family 1.7
Nedd4-2: Neuronal precursor cell expressed developmentally downregulated-4 type 2.
SNI: Spinal nerve injury
TTX-S: Tetrodotoxin-sensitive
V1/2: Midpoint potentials

## Acknowledgements

Funding: Supported by NINDS [NS098772 and NS120663 (R.K.)], NIDA (DA042852, R.K.) Author contributions: R.K. and A.M. developed the concept and designed experiments; A.M., K.G., and D.R., collected and analyzed data; C.L.M. performed animal behavior studies; R.K. provided funding; D.R., K.G., R.K. and A.M. wrote the manuscript; and R.K. and A.M. supervised all aspects of this project. All authors had the opportunity to discuss results and comment on the manuscript; Data and materials availability: All data is available in the main text and figures.

